# ERAASR: An algorithm for removing electrical stimulation artifacts from multielectrode array recordings

**DOI:** 10.1101/185850

**Authors:** Daniel J. O’Shea, Krishna V. Shenoy

## Abstract

Electrical stimulation is a widely used and effective tool in systems neuroscience, neural prosthetics, and clinical neurostimulation. However, electrical artifacts evoked by stimulation significantly complicate the detection of spiking activity on nearby recording electrodes. Here, we present ERAASR: an algorithm for Estimation and Removal of Artifacts on Arrays via Sequential principal components Regression. This approach leverages the similar structure of artifact transients, but not spiking activity, across simultaneously recorded channels on the array, across pulses within a train, and across trials. The effectiveness of the algorithm is demonstrated in macaque dorsal premotor cortex using acute linear multielectrode array recordings and single electrode stimulation. Large electrical artifacts appeared on all channels during stimulation. After application of ERAASR, the cleaned signals were quiescent on channels with no spontaneous spiking activity, whereas spontaneously active channels exhibited evoked spikes which closely resembled spontaneously occurring spiking waveforms. The ERAASR algorithm requires no special hardware and comprises sequential application of straightforward linear methods with intuitive parameters. Enabling simultaneous electrical stimulation and multielectrode array recording can help elucidate the causal links between neural activity and cognitive functions and enable the design and implementation of novel sensory protheses.

## 1 Introduction

Electrical stimulation is a method of modulating neural activity that is widely used within neuroscience and neuroengineering, as well as for treatment of chronic neurological pathologies. Within neuroscience, microstimulation is used to probe the functional organization of neural circuits. Whereas electrophysiological recordings provide correlative insights that neural activity in a brain region appears related to a specific behavior, electrical stimulation can demonstrate causal contributions of brain regions to specific cognitive functions (Salzman et al., 1990). In particular, intracortical microstimulation (ICMS) has been used to identify neural networks underlying perception, attention, cognition, and movement (see Cohen and Newsome, 2004; Clark et al., 2011; Histed et al., 2013, for a review). ICMS can also drive reliable percepts in humans and animal models, including somatosensory vibration (Romo et al., 2000) and visual phosphenes (Tehovnik and Slocum, 2007), which underscores its application for delivering artificial sensation in visual (Tehovnik et al., 2009) and motor prosthetic systems (Berg et al., 2013; O’Doherty et al., 2011). In clinical contexts, stimulation, especially delivered to deeper brain structures, has demonstrated efficacy for the treatment of neurological and neuropsychiatric conditions (Wichmann and Delong, 2006; Holtzheimer and Mayberg, 2011) and recently as an avenue for providing sensory information as will be needed for a range of emerging brain-machine interface systems (Flesher et al., 2016).

However, as with all perturbations to the nervous system, accurate interpration of the results relies on a solid understanding of the effects of the perturbation made (e.g., Jazayeri and Afraz, 2017; Otchy et al., 2015). With lesion and pharmacology studies, understanding the size of the affected region, completeness of the intended effect, the timecourse of the effect and recovery, and the influence of compensatory or homeostatic adaptation mechanisms are all critical to drawing correct conclusions from the behavioral impairments observed. Analogously, with ICMS, it is essential to understand the effect that electrical stimulation has on neural activity in order to draw correct inferences from causal experiments. In the context of neural prosthetics, researchers aim to improve the fidelity of artificial perception with sophisticated spatiotemporal patterning (e.g., Dadarlat et al., 2015) and to optimize stimulation paramaters to maximize therapeutic benefit. Obtaining a clear picture of the effects of particular stimulation paradigms will likely prove essential to this goal, enabling closed-loop tuning of stimulation parameters relative to a desired effect on the local neural activity.

The effects on neural firing patterns evoked by ICMS have been difficult to characterize because the electrical artifact induced by stimulation interferes with the electrophysiological recording equipment used to record neuronal responses (Ranck, 1975; Merrill et al., 2005). A variety of approaches have been developed to work around these artifacts, which can be organized into several categories. Online approaches employ special hardware within the signal recording pathway to remove the stimulation artifact online (e.g., Wichmann and Devergnas, 2011; Brown et al., 2008; Müller et al., 2012). While effective, hardware approaches can be expensive or challenging to scale to high channel count recording arrays and typically require very stable artifacts over time for proper removal. Offline approaches employ standard electrophysiology collection hardware to record neural signals and then estimate and subtract the artifact post hoc. The most common approach for artifact estimation is to simply interpolate across the artifact (O’Keefe et al., 2001; Montgomery et al., 2005) and/or to average over repeated stimulation epochs (Hashimoto et al., 2002; Montgomery et al., 2005; Klink et al., 2017) and subtract, exploiting the larger stereotypy of the stimulus artifact relative to variable neural responses. Others have employed curve fitting approaches to estimate the artifact, exploiting the differences in shape of artifact and action potentials, thereby allowing for small variations in the observed artifact across trials (Wagenaar and Potter, 2002; Erez et al., 2010).

These previous approaches have two major shortcomings addressed in this work. First, these approaches only recover neural signals occurring after the stimulation pulse is completed, with a typical window of of several milliseconds during which signal is discarded. However, ICMS is typically delivered in trains of stimulation pulses occurring at frequencies approaching several hundred pulses per second (e.g., Churchland and Shenoy, 2007; Klink et al., 2017). This failure to recover neural activity concomitant with the stimulation pulses renders a large fraction of the peri-stimulation signal unusable for spike detection and waveform discrimination. Secondly, prior work has focused on artifacts detected at a single electrode, not exploiting the similar structure of contaminating artifacts across multiple electrodes using a multielectrode array. A notable exception to both of these shortcomings was developed by Mena and colleagues (2016), where they addressed both of these limitations using a modern statistical framework to isolate electrical artifacts from neural spiking signals in multielectrode array recordings of the retina *in vitro*. However, this approach currently requires the availability of electrical images for each neuron’s spike waveform on multiple surrounding electrodes. Unfortunately, for the multielectrode array configurations commonly employed in *in vivo* cortical electrophysiology, the electrical image of most neurons is confined to a single electrode. Additionally, since the set of recorded neurons is biased towards a highly active subset of neurons (Barth and Poulet, 2012), we anticipate that some neurons that are driven to spike by electrical stimulation may be quiet outside of the stimulation period, and thus ommitted in the dictionary of available electrical images.

In this paper, we present a fast method for removing electrical stimulation artifacts from multielectrode array recordings. We term this approach ERAASR: Estimation and Removal of Artifacts from multielectrode Arrays via Sequential principal components Regression. ERAASR does not require any special hardware and can recover the full timecourse of neural signals during and after stimulation, requiring only that the electrical artifact does not saturate the amplifier on the recording electrodes. Our algorithm exploits the similarity of artifacts across multiple electrodes, multiple pulses within a pulse train, and multiple trials of repeated stimulation. The ERAASR algorithm is structured as a sequence of cascaded, linear operations—PCA and linear regression—across each axis of the data tensor. We validate our method on neural signals collected during an arm-movement task with 24-channel linear multielectrode arrays (“V probes”, Plexon Inc.) in macaque dorsal premotor cortex, demonstrating that spiking waveforms extracted during the stimulation period closely resemble spontaneous spiking waveforms on each electrode.

## 2 Methods

### 2.1 Subjects

Animal protocols were approved by the Stanford University Institutional Animal Care and Use Committee. The subject was one adult male macaque monkey (*Macaca mulatta*), monkey P. After initial training, we performed a sterile surgery during which the macaque was implanted with a head restraint and a recording cylinder (NAN Instruments) which was located over left, caudal, dorsal premotor cortex (PMd). The cylinder was placed surface normal to the skull and secured with methyl methacrylate. A thin layer of methyl was also deposited atop the intact, exposed skull within the chamber. Before stimulation sessions began, a small craniotomy (3 mm diameter) was made under ketamine/xylazine anesthesia, targeting an area in PMd which responded during movements and palpation of the upper arm (17 mm anterior to interaural stereotaxic zero).

### 2.2 Reaching task

For the purposes of this study, we engaged the monkey in a reaching task in order to drive task related neural activity in premotor cortex where our electrodes were located. We trained the monkey to use his right hand to grasp and translate a custom 3D printed handle (Shapeways, Inc.) attached to a haptic feedback device (Delta.3, Force Dimension, Inc.). The other arm was comfortably restrained at the monkey’s side. The monkey was trained to perform a delayed reaching task by moving the haptic device cursor towards green rectangular targets displayed on the screen. Successful completion of each movement triggered a juice reward.

### 2.3 Electrophysiology and stimulation configuration

At the start of each experimental session, both a stimulation electrode and a recording probe were secured to two independently controllable, motorized micromanipulators (NAN instruments). Both probes were lowered simultaneously through blunt, non-penetrating guide tubes into dorsal premotor cortex at 3 μm/s to an approximate depth of 2 mm (Figure 1a). The stimulation electrode was a single tungsten microelectrode with approximately 1MΩ impedance at 1 kHz (Part #UEWLGC-SECN1E Frederick Haer Company). The recording probe was a linear electrode array consisting of 24 contact sites located at 100 μm spacing along the length of the shank (V-probe PLX-VP-24-15ED-100-SE-100-25(640)-CT-500, Plexon Inc.). The recording probe penetration site was located at an approximate distance of 0.75mm to 1.5mm from the stimulation site. As best possible, the stimulating electrode was inserted at the same location and to the same depth, whereas we changed the location of the recording electrode for each recording session.

**Figure 1.**
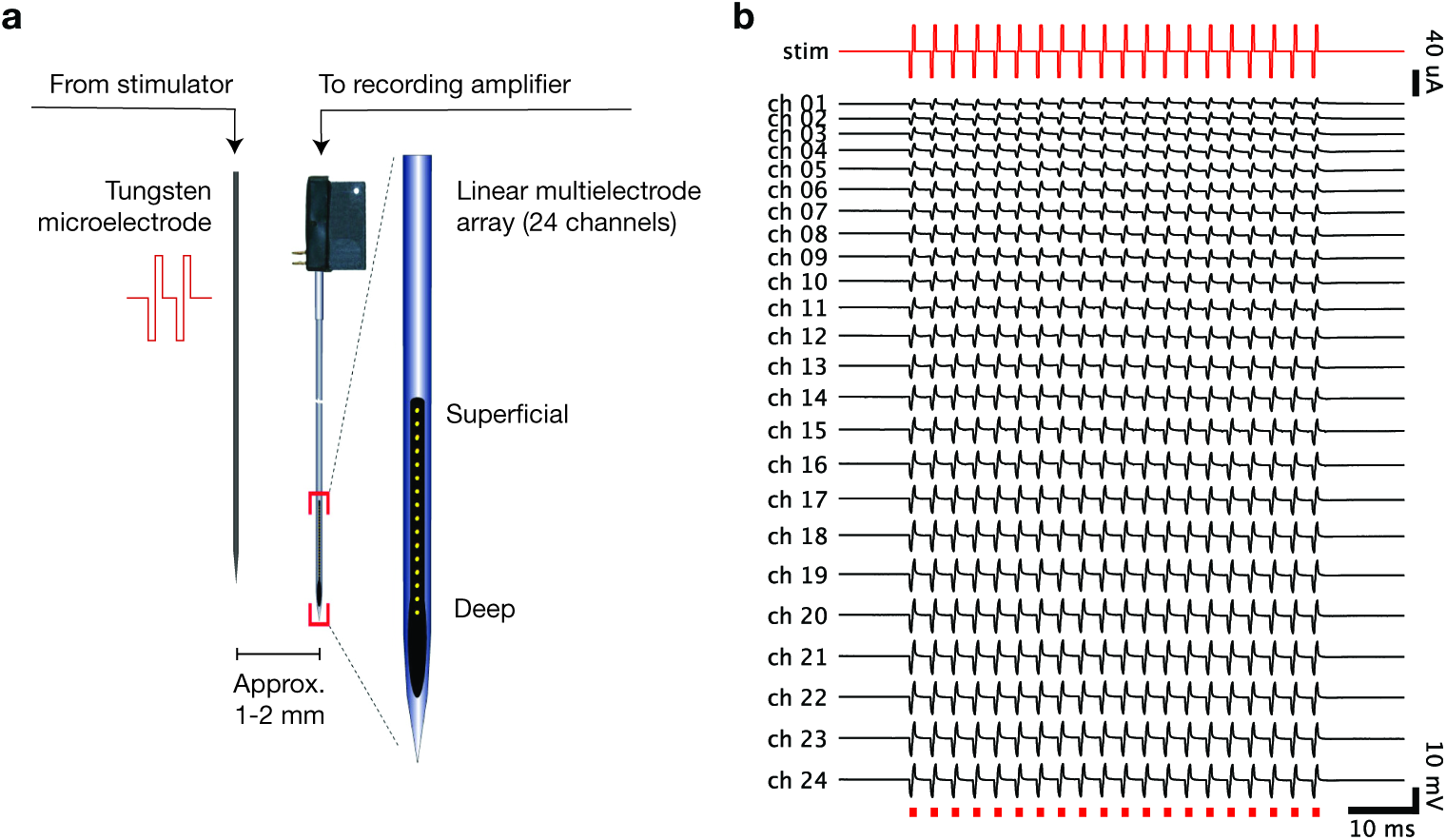
Our technique faciliates electrical recordings of local neural activity during and after nearby electrical stimulation, provided that the electrical artifact does not saturate the amplifier. (a) A schematic of the recording setup. A tungsten stimulating microelectrode and linear multielectrode array are inserted in parallel into dorsal premo-tor cortex. The linear array records the electrical activity of nearby neurons spanning the layers of cortex, while electrical current is introduced by the tungsten microelectrode (b) An example of real electrical recordings on the linear multielectrode array (ch. 1-24 during delivery of a single 40 μApulse train (red, stim) through the stimulating electrode The recording channels span 2.3 mm and are labeled from 1 (most superficial, typically above cortex) through 24 (deep, typically in white matter).

The stimulation was performed using a StimPulse electrical stimulation system used as a combined function generator and isolated current source (Frederick Haer Company). The electrical stimulation current flowed through the brain via the electrode to a “ground screw” located at the posterior pole of the implant whose tip was in contact with the dura below the skull. Microstimulation trains consisted of 20 biphasic pulses delivered at 333 Hz for 57 ms (150 μs cathodic, 100 μs pause, 150 us anodic). Stimulation amplitude was varied between 5 μA to 40μA. Stimulation delivered at the start of most experimental sessions, while the arm was in a passive, resting position, triggered brief movements of the contralateral upper arm and shoulder at thresholds typically *>* 140 μA. Stimulation onset was triggered via TTL pulse delivered by the task engine (Simulink Real-time Target, The Mathworks; NIDAQ digital to analog card National Instruments). Stimulation was delivered on 20-40% of trials and randomly, interleaved. Stimulation amplitude was fixed within each block of trials, where we collected at least 8-10 repetitions of each stimulation trial type for each block. Within each session, we typically began with blocks of lower amplitude stimulation before proceeding to higher amplitude stimulation.

The recording probe was connected to a 3-headed switching headstage, a component of the Blackrock StimSwitch system (Blackrock Microsystems. The probe connects to one bank of inputs, control lines from the StimSwitch control box connect to the second bank of inputs, and outgoing voltage signals leave via the third bank of outputs. During these experiments, this component was configured to act as a unity gain buffer (head stage) which simply relays *the* voltage signals to a shielded ribbon cable (Samtec Inc.) to the amplifier (Blackrock Microsystems). Each electrode is differentially amplified relative to a common reference line in the V-probe itself, which is also shorted to surrounding guide tube for better noise rejection.

Broadband voltages were recorded from all 24 electrode channels of the recording probe. We also recorded from the stimulation electrode during initial penetration to verify that we could see nearby neurons on the electrode. We then disconnected the stimulation electrode from the recording system and connected it to the StimPulse stimulator. Broadband signals were filtered at the amplifier (0.3 Hz 1 pole high-pass filter, 7.5 kHz 3 pole low-pass filter), digitized to 16 bit resolution over ± 8.196 mV (resolution = 0.25 μV), and sampled at 30kHz.

### 2.4 Artifact amplitude model

The amplitude of the electrical artifact was measured on each recording channel as the peak to peak voltage concomitant with the first stimulus pulse, averaged across trials. These amplitude measurements were used to fit a model relating artifact amplitude to the distance of each site to the stimulation source. This model jointly optimizes a single scaling parameter for the distance-dependent amplitude relationship as well as parameters denoting the stimulation-relative coordinates of the recording probe in each session. We define *x*_*s*_ as the distance measured laterally from the probe to the closest point on the stimulating electrode, both of which are assumed to run parallel to the recording probe and normal to the cortical surface; we define *y*_*s*_ as the vertical distance along the penetration path from the most superficial electrode on the probe to the tip of the stimulating electrode. We begin by describing the location of each electrode *e* in each session *s* relative to the probe location (*x, y*)_*s*_. Here

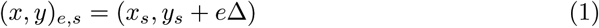

where Δ = 100 μm spacing between sites. This contact site is located at a distance

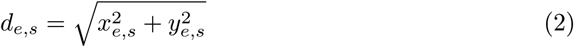

from the stimulation source defined as the origin. We then assume that the amplitude of the artifact on each electrode for a specific stimulation amplitude *I*_*a*_, falls of as the reciprocal of this distance.

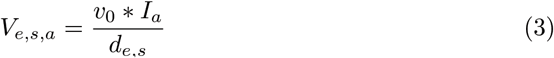

where *v*_0_ has units V/(μA mm). The parametersof thie model are this single scaling parameter *v*_0_ as well as a pair of location coordinates for the probe on each session.

We then fit this model using constrained non-linear curve fitting (lsqcurvefit in Matlab) to minimize the squared error between empirical and modeled artifact amplitude across electrodes, sessions, and amplitudes. Confidence intervals of the parameters were obtained using the Jacobian of the fitted model.

### 2.5 Precise temporal alignment and tensor construction

We begin with the raw, unfiltered voltage traces for each stimulation trial, collected simultaneously on the 24 channels, sampled at 30 kilosamples/sec. We first correct the small amount of jitter between triggering the stimulator to the beginning of the stimulation train, typically on the order of a few milliseconds. We begin by precisely aligning the stimulation trials to each other in time. This process is performed separately for each group of trials sharing a common stimulation amplitude. We extract the raw voltage data off of one electrode channel in a 60 ms window surrounding the triggering event, high-pass filter the signal with a fourth-order, 250 Hz corner frequency Butterworth filter, to remove slow drifit. We then detect the onset of the artifact due to the first biphasic pulse on each trial using an appropriately low negative threshold, set to avoid spiking activity but reliably detect artifacts at the lowest stimulation amplitude. We then realigned the voltage data for each trial to this threshold crossing event. We then performed an outlier detection procedure to detect rare instances of stimulator failure, including trials where 19 or 21 pulses were delivered instead of 20. Outlier detection was performed by projecting the realigned, unfiltered voltage traces onto the first 10 principal components of that group, and detecting trials where the absolute value of the z-scored scores in any of the first 10 PCs exceeded a value of 5. Outlier trials (typically *<* 2% of trials) were excluded from subsequent analysis. The traces were then up-sampled 10x to 300 kHz using spline interpolation and then a maximum cross-correlation procedure was performed to determine the temporal offset between each trial and the first in the group. We manually verified the results of this procedure to ensure successful artifact removal and accurate alignment of the artifact pulse trains. The aligned traces are then resampled at the original 30 kHz sampling rate.

The ERAASR algorithm is summarized in alg. 1. We construct a data tensor for each group of trials sharing a common stimulation amplitude, as the artifact evoked across trials for a single stimulation amplitude were highly similar. We extract a 60 ms window of the trial-aligned broadband voltage data across the trials, yielding a data tensor X_raw_ with size *C* (number of channels) x *T* (number of timepoints per pulse) x *P* (number of pulses) x *R* (number of trials). In our datasets *C* = 24 channels, *T* = 90 timepoints per pulse (3 ms at 30 kHz sampling), *P* = 20 pulses, and *R* was typically on the order or 100-200 trials.

We lightly high pass filter the signals in time with a fourth-order, 10 Hz corner frequency Butterworth filter, which removes slow drifits from the traces and makes subsequent processing more robust. We first attempt to clean each channel by exploiting the similarities of the artifact across simultaneous channels (Figure 2a).

**Figure 2.**
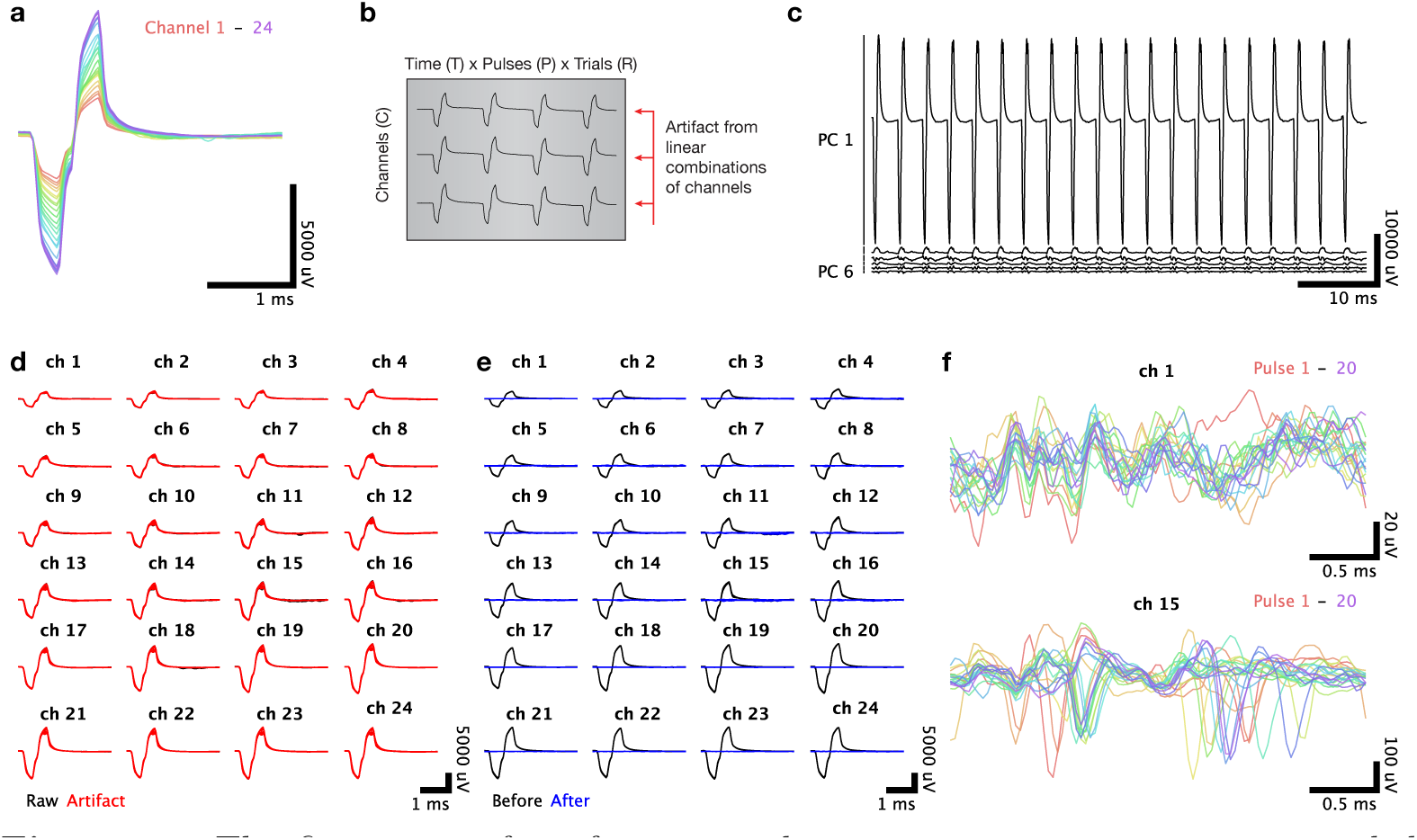
The first stage of artifact removal removes common structure recorded simultaneously across linear electrode array channels. (a) The electrical artifact evoked by a single pulse of a stimulus train is highly similar across the 24 channels of the array. (b) To identify the common artifact structure present across channels, we rearrange the artifact data into a matrix with dimensions time × pulses × trials (*T × P × R*) by channels (*C*), shown transposed. (c) The principal components of this matrix are linear combinations of the electrode channels. The projections of the recordings along these components clearly display artifact structure shared across channels. (d) Voltage recordings for the first pulse of the stimulus train, plotted separately for each channel with the responses on each trial superimposed. Raw voltage recordings are shown in black, with the artifact inferred via principal components regression shown in red. (e) Same presentation as in (d) but with the residual voltage recordings shown in blue, afiter the inferred artifact has been subtracted. (f) Cleaned voltage recordings for a single trial with the responses to each individual pulse superimposed. The top panel shows a superficial channel which had no detectable spontaneous neural activity, and the bottom panel shows a lower channel where a well-isolated single unit was spiking spontaneously.

### 2.6 Removal of common structure over channels

We then unfold the data tensor into an *RTP* x *C* matrix M^*c*^ (Figure 2b), and perform principal components analysis. This allows us to re-express each channel’s response vector (*RTP* x 1) as a linear combination of principal components, each *RTP* x 1. As the artifact waveform is very large relative to the interesting spiking activity and shared across all channels, one would expect the top few principal components to capture preferentially shared variance due to the artifact. Empirically, we determined that the top *K*_*C*_ = 4 principal components (PCs) captured much of the artifact shape (Figure 2c). We then want to reconstruct each channel’s response omitting the contribution from the artifact. The simplest method to accomplish this would be to reconstruct the channel’s response by using a linear combination of all PCs except the first *K*_*C*_. However, in some cases, especially when lower currents were used and the magnitude of the artifact relative to the spiking activity was smaller, we observed that some of the top PCs would begin to incorporate a small amount of spiking activity from individual channels. By reconstructing those channels from the remaining PCs, the spiking activity itself would be partially distorted because a fractional portion of those spikes would be subtracted along with the artifact. We addressed this issue by again exploiting the locality of spiking activity, by assuming that spiking activity from any one channel would not be present on other channels except the immediately adjacent channels. However, the artifact is shared on a much larger spatial scale and can be separated from the spiking activity. Therefore, for each channel *c*, we used the following procedure: reconstruct the top *K*_*c*_ PCs using all channels except *c* and its immediately adjacent neighbors *c* − 1 and *c* + 1, that is, using a modified version of the loading weights 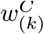 for the *k*th principal component in which we set the weights for *c, c* − 1, *c* + 1 to 0. (More generally, we define parameter λ_*C*_ that dictates the number of adjacent channels excluded from the reconstruction. We refer to this vector of modified loading weights as 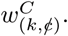

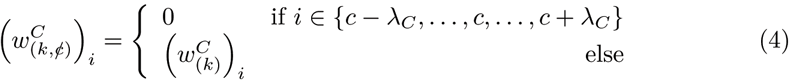

#### Algorithm 1

ERAASR cleaning procedure

**Input**: Raw data tensor X_raw_(*c,t,p,r*) with size *C* channels by *T* timepoints per pulse by *P* pulses by *R* trials

**Output**: Cleaned data tensor X_cleaned_(*c,t,p,r*) with same size

**Parameters**: *K*_*C*_, *K*_*P*_, *K*_*R*_ - number of principal components to describe artifact structure over channels, pulses, trials

λ_*C*_, λ_*P*_, λ_*R*_ - the number of adjacent channels, pulses, trials excluded as regressors during artifact reconstruction *β*_*P*_, *β*_*C*_ ∈ {*false true*} - Perform cleaning over pulses separately on each channel? Below we assume *β*_*P*_ is false and *β*_*C*_ is true for clarity.

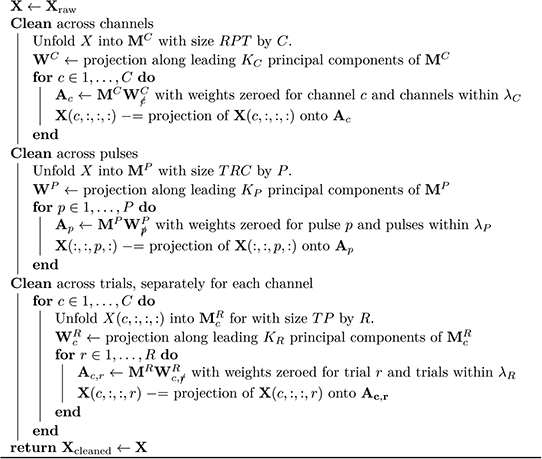

Using these loading weights, we reconstruct the matrix of the top *K*_*C*_ artifact-capturing PCs where each column is given by

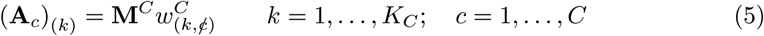

We then regress the channel’s response vector against these artifact components and subtract this reconstructed artifact from the channel’s response:

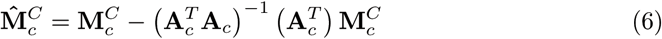

This cleaning across channels captured (Figure 2d) and removed (Figure 2e) much of the artifact from each pulse. However, when examining the responses on an individual channel across each of the 20 pulses superimposed, we observed consistent structure in the traces (Figure 2f).

### 2.7 Removal of common structure over pulses

This structure indicates that there remained artifact structure that is unique to individual channels but shared among the pulses in each trial. To address this, we rearrange 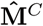 into a new matrix M^*P*^ with size *TRC* x *P*, a set of responses to each individual pulse, concatenated over trials and channels (Figure 3a). Empirically, we determined that the top *K*_*P*_ = 2 principal components PCs) captured much of the artifact shape. Consequently, we used the same PC regression procedure to estimate and subtract the artifact on each pulse, omitting the contribution of the pulse under consideration and optionally adjacent pulses within *λ*_*P*_. For our dataset, *λ*_*P*_ = 0 was used. If 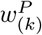 is a vector of *P* loading weights describing the *k*th principal component, and for each pulse *p*, 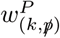 is given by:

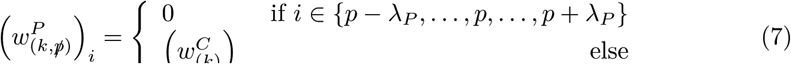

**Figure 3.**
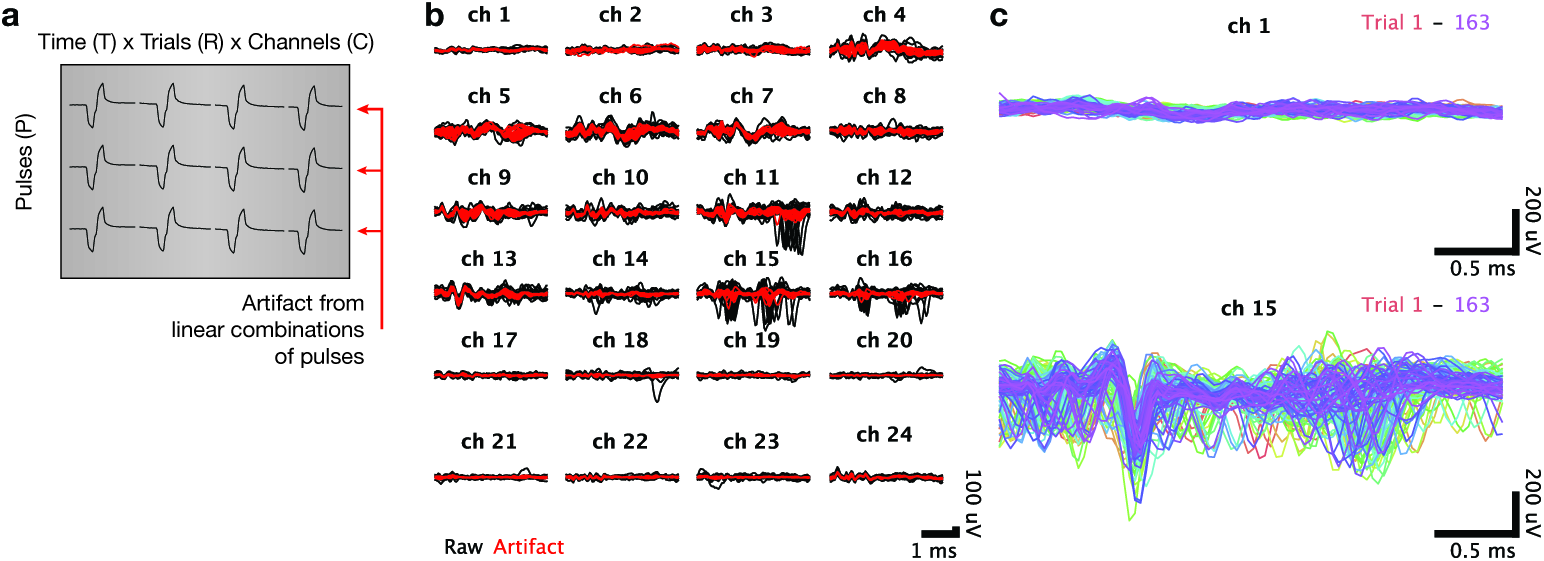
The second stage of artifact removal removes common artifact structure across the successive pulses of the stimulus train. (a) To isolate structure across pulses, we rearrange the artifact data into individual matrices for each channel, each with dimensions time (*T*) × trials *(R)* × channels (*C*) by pulses (*P*). (b) Voltage recordings for a single trial, plotted separately for each channel with the responses to each pulse superimposed. Raw voltage recordings are shown in black, with the artifact inferred via principal components regression shown in red. (c) Cleaned voltage recordings for a single pulse with the responses to all trials superimposed. The top panel shows a superficial channel which had no detectable spontaneous neural activity and the bottom panel shows a lower channel where a well-isolated single unit was spiking spontaneously.

For each pulse, we reconstruct the matrix of the top *K*_*P*_ artifact-capturing PCs, whose columns are given by:

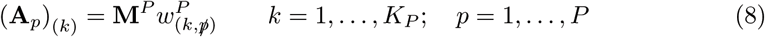

We then regress the channel’s response vector against these artifact components and subtract this reconstructed artifact from the channel’s response:

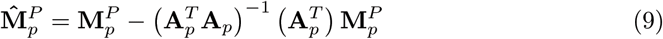

This cleaning procedure over pulses captured much of the common artifact structure observed within each train (Figure 3b) It is also possible to perform this cleaning over pulses separately for each channel, following an approach like that described below for cleaning over trials.

We next examined the set of cleaned traces for individual channels and pulses in the train, over the full set of trials. We observed that there remained common artifactual structure in these traces, which suggests that there is artifact structure unique to that channel and pulse number but shared among many trials (Figure 3c). In most cases, this common structure was similar among nearby trials but exhibited a slow drifit in the shape of the artifacts over the experimental session.

### 2.8 Removal of common structure across trials

To remove these residual artifacts, we again employed a principal components regression approach, exploiting the shared structure of the artifact over multiple trials. We performed this operation separately for each channel. We rearranged the cleaned 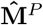 into a set of *C* data matrices 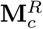 for *c* ∈ {1,…, *C*}, each with size *TP* x *R* (Figure 4a). Empirically, we determined that the top *K*_*R*_ = 4 PCs captured much of the artifact shape over trials. If 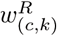 is a vector of *R* loading weights describing the *k*th principal component for channel *c*, we defined:

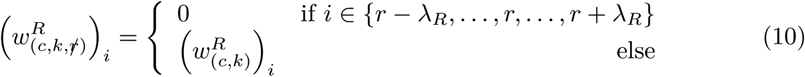

where *λ*_*R*_ adjacent trials were excluded from the reconstruction. For our dataset, *λ*_*R*_ = 0 was used.

**Figure 4.**
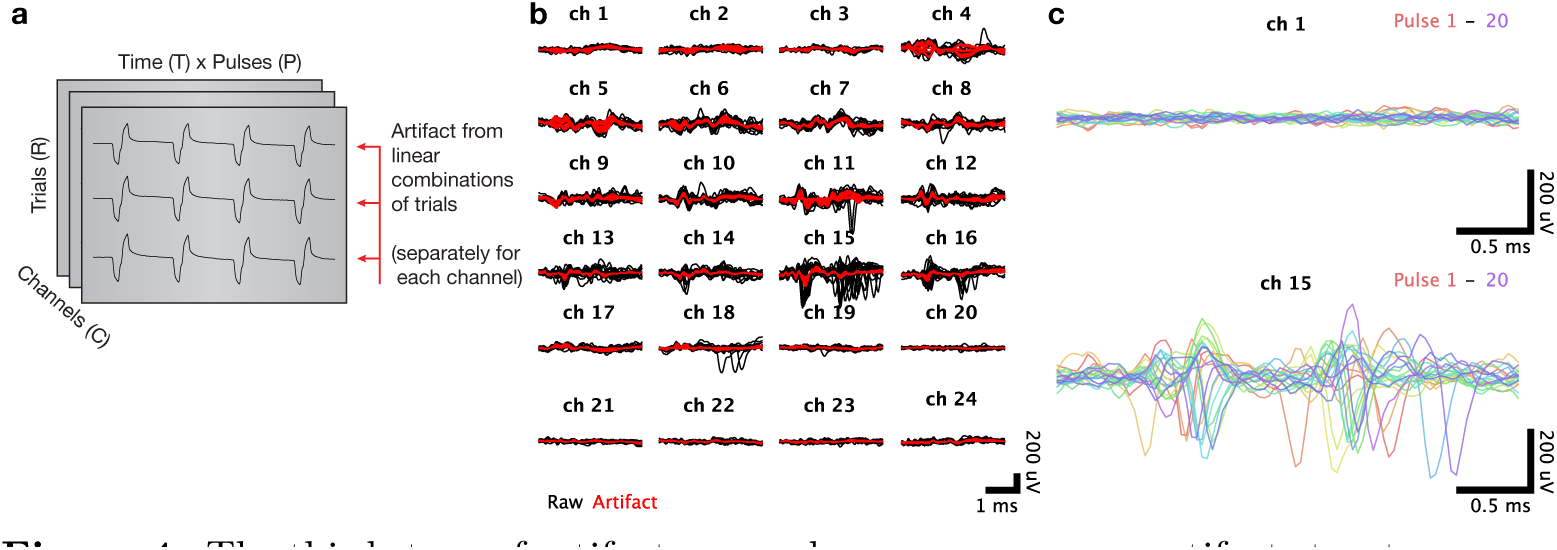
The third stage of artifact removal removes common artifact structure across trials in which identical electrical stimulus trains were delivered. (a) To isolate structure across trials, we rearrange the artifact data into individual matrices for each channel, each with dimensions trials *(R)* by time × pulses *(T × P)*. (b) Voltage recordings for a single pulse, plotted separately for each channel with the responses to each trial superimposed. Raw voltage recordings are shown in black, with the artifact inferred via principal components regression shown in red. (c) Cleaned voltage recordings for a single trial with the responses to all pulses superimposed. The top panel shows a superficial channel which had no detectable spontaneous neural activity and the bottom panel shows a lower channel where a well-isolated single unit was spiking spontaneously.

For each trial, we reconstruct the matrix of the top *K*_*R*_ artifact-capturing PCs, whose columns are given by:

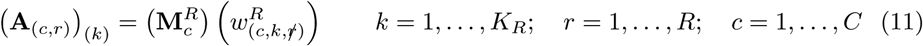

We then regress the trial’s response vector against these artifact components and subtract this reconstructed artifact from the trial’s response:

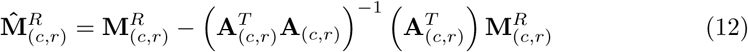

We then rearranged the set of cleaned 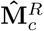 back into a *C* x *T* x *P* x *R* tensor X_cleaned_. This cleaning procedure captured much of the residual artifact (Figure 4c). The cleaned signals in X_cleaned_ did not display features obviously indicative of residual stimulation artifact, but did on some channels contain readily detectable spiking activity (Figure 4c, bottom panel). We then reinserted X_cleaned_ into the raw voltage traces for each stimulation trial, using appropriate offsets to preserve continuity at the first and last timepoints of the inserted segment.

### 2.9 Post-stimulation transient cleaning

Following stimulation offset, a slower transient was observed in all channels. These transients are highly similar among trials. Consequently, we employed a similar principal components regression procedure to remove these slow transients. We performed the following procedure for each stimulation amplitude separately. We began by aligning the trials to stimulation offset and extracting voltage data in a 30 ms time window beginning afiter stimulation end, yielding a tensor X_post_ with size *R* by *T* by *C*. We then rearrange this tensor into a set of *C* data matrices 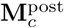 for *c* ∈ {1,…, *C*}, each with size *T* x *R*. Empirically, we determined that much of the transients temporal structure was present in each trial for a given channel and that the top *K*_post_ = 2 principal components (PCs) of these 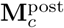 matrices captured much of the transient shape over trials Consequently, we repeated the same principal components regression approach described in eqs. (10) to (12) to clean the post stimulation transients over trials. We then rearranged the set of cleaned 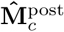 back into a *R* by *T* by *C* tensor X_post-cleaned_. The signals in X_post-cleaned_ were then reinserted into the full voltage traces for each stimulation trial.

### 2.10 Spike thresholding and sorting

The cleaned voltage traces, along with the original voltage traces on non-stimulated trials were then high-pass filtered using a fourth-order 250 Hz corner frequency Butterworth filter, as is typically done online before spike detection. We then thresholded each signal at −4.5x RMS voltage, using a greedy procedure in which the spike threshold crossings were detected in order of size (largest to smallest), and no further threshold crossings were considered within a lockout period extending 0.3 ms prior to and 1.0 ms afiter the current threshold crossing. We then hand-sorted the spiking waveforms using MKsort (Matthew Kaufman and Ripple, Inc., https://github.com/ripple-neuro/mksort).

### 2.11 Unit selection

During experiments, we recorded any units or multi-units that could be isolated on the multielectrode array, without regard for their responsiveness or modulation by the task. Before analysis, we applied a screening procedure which looked only at non-stimulated trials in order to remove neurons that were very unreliable or very weakly modulated by the task. Briefly, we filtered non-stimulated trials spike trains with a 30 ms Gaussian window, then aligned trials separately from 100 ms pre-target onset to 70 ms post go cue, and then from 300 ms pre-movement to 600 ms post, then averaged within groups of trials with the same reach target. We defined each unit’s “signal” to be the range of firing rates over all times and conditions, and “noise” to be the maximum standard error of the mean firing rate. We included units where the ratio of signal to noise exceeded at least 2, a total of 138 units in monkey P. Note that this selection process looked only at non-stimulation trials.

## 3 Results

We delivered high-frequency (333 Hz) biphasic electrical stimulation in macaque dorsal premotor cortex through a tungsten microelectrode, while simultaneously recording neural spiking activity using a second 24-channel linear multielectrode array (Figure 1a). When a stimulus train was delivered through the electrode, a highly stereotyped stimulation artifact appeared on all channels of the recording array (Figure 1b). These broadband recordings were corrupted by a very large electrical stimulation artifact. Amplitudes could exceed ±7500 μV which is nearly two orders of magnitude larger than spiking waveform amplitudes (~100 μV). The artifact traces resembled filtered versions of the original current stimulus. with similar transients evoked for each of the 20 biphasic pulses. These artifacts increased in amplitude with larger stimulation current and smoothly varied over the 24 channels of the recording probes (Figure 5a,b), consistent with a distance-dependent attenuation of the electrical stimulus.

We used these amplitude profiles over channels for each session to fit a model relating artifact amplitude and the reciprocal of the distance of each electrode to the stimulation source. The model fitting process jointly optimizes a scaling parameter of the amplitude-distance relationship as well as parameters specifying the location of the recording probe on each session (See methods). We fit this model to data collected in 9 recording and stimulation sessions in macaque prefrontal cortex. Figure 5c shows the empirical and predicted artifact amplitude (normalized by stimulation current) under the model against distance from the stimulation site (squared correlation *R*^2^ = 0.91), demonstrating that this model well-describes the distant-dependent falloff of the electrical artifact. The fitted location of each probe relative to the stimulation source are depicted in Figure 5d, which closely aligns with the noted approximate probe insertion locations at 0.7mm to 2mm from the recording probe. This simple relationship suggests that under certain circumstances, electrical artifacts could be used to reconstruct recording locations *in vivo*.

**Figure 5.**
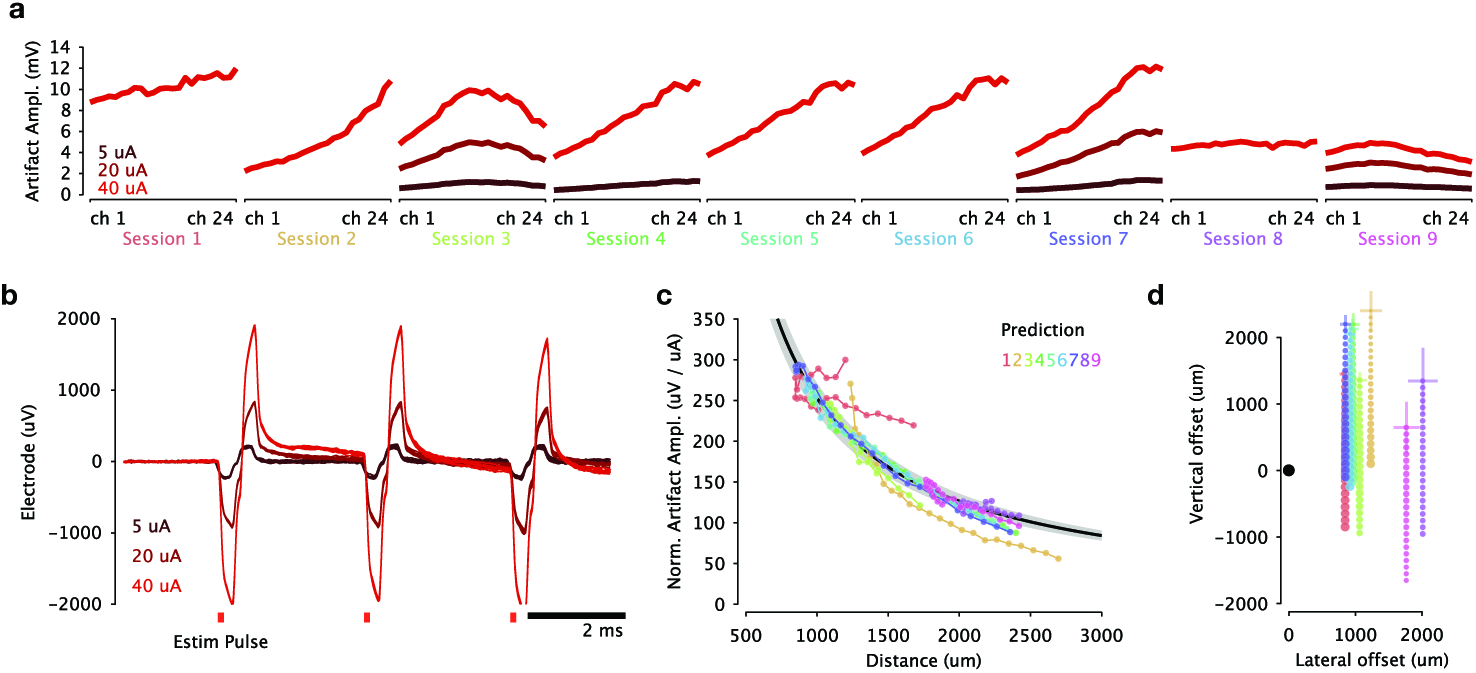
Electrical artifacts recorded on the electrode array resemble scaled, filtered versions of the original stimulus pulse train. The artifact amplitude varies smoothly with stimulation current and electrode distance, enabling *post hoc* inference of the locations of the recording probes in each session. (a) Electrical artifact amplitude (peak to peak) across multielectrode array channels. Each plot corresponds to a recording session in which the multielectrode array was inserted in a new location. (b) Electrical artifacts recorded on several trials on superficial electrode with increasing electrical stimulation current. (c) A simple model relating the electrical artifact amplitude was fit to each channels on every session. The model assumes that the artifact scales linearly with stimulation current and falls off with the reciprocal of Euclidean distance from the stimulating electrode tip. These distances are themselves inferred from the data as parameters describing the position of the electrode array in each session. The fitted model predicted amplitudes (black curve flanked by gray 95% CI) align well with the empirical amplitudes observed on each session (groups of connected dots, each color corresponds to a session). (d) The fitted locations of the recording probe on each session relative to the stimulating electrode tip. Marker size is scaled according to artifact amplitude. Cross-hairs indicate 95 % CI for probe location parameters.

Having characterized the artifacts, we applied ERAASR to remove the artifact from these traces. The ERAASR algorithm leverages several features of these recording datasets. First, it assumes that the true neural signal is corrupted by an additive stimulation artifact, which depends critically on the assumption that no information is lost due to amplifier saturation during the stimulation period. For the 5 μA to 40 μA range of stimulation amplitudes used, the stimulation artifact produced did not saturate the amplifier’s range (±8196μV) on any of the recording channels in any experimental session. Therefore, if the artifact shape can be properly estimated, it can be removed via simple subtraction. Second, because we recorded simultaneously on multiple, closely spaced channels, we have multiple simultaneous observations of the electrical stimulation artifact at different points in space, whereas the individual channels do not share the same spiking waveforms, especially if we exclude immediately adjacent channels. Third, the stimulation train contains 20 repeats of identical biphasic pulses, which produced highly similar artifact transients. Finally, we deliver the stimulation train over many trials, yielding similar artifacts on each repetition. Because spiking activity evoked during the stimultion period is unlikely to occur at precisely the same time relative to each pulse and on each trial, we can use these repeated artifact measurements to build an estimate of the artifact shape for subsequent removal.

**Figure 6.**
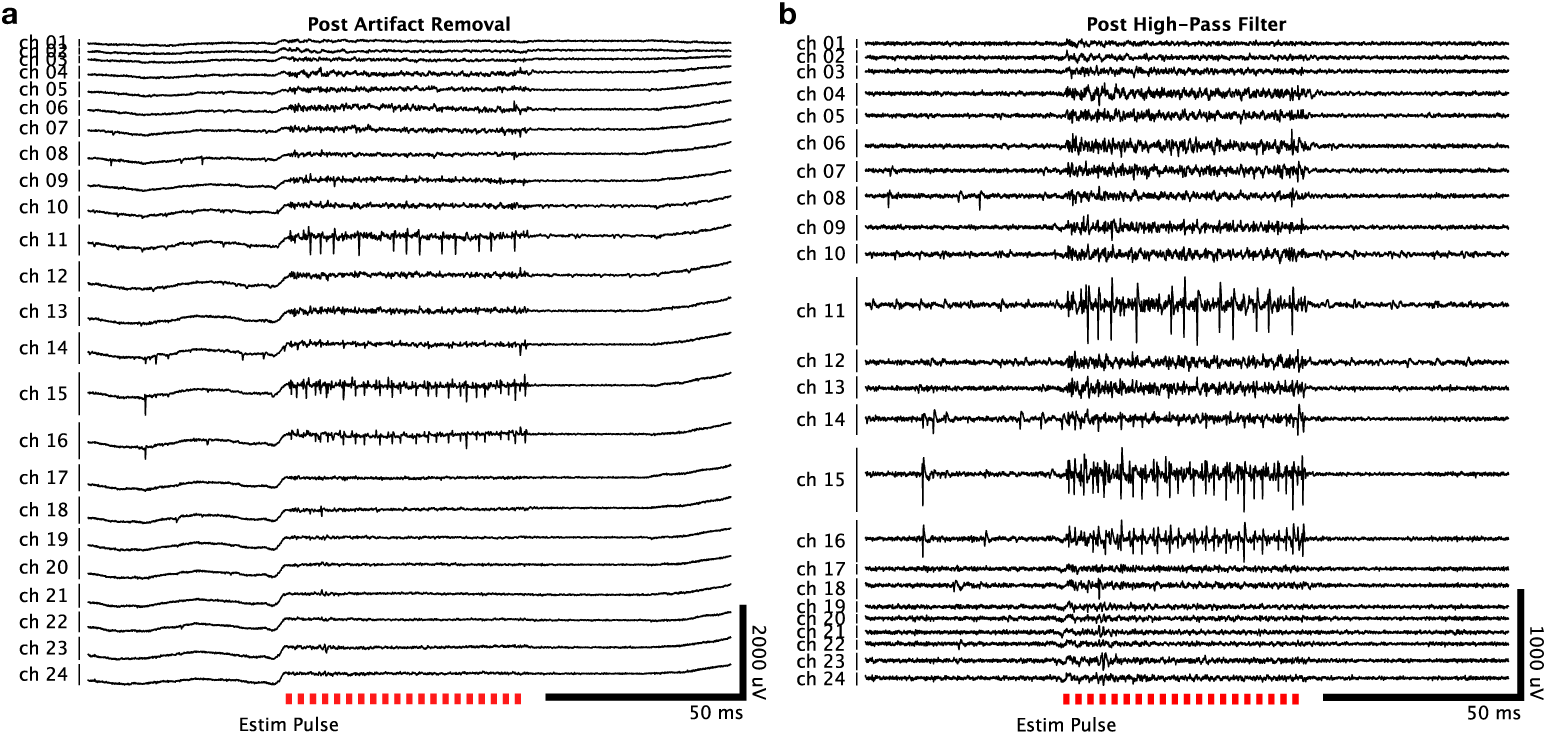
The cleaned single trial voltage recordings exhibit very little residual stimulation artifact. (a) Electrical recordings on the linear multielectrode array for the same trial shown in 1(b). Spiking activity during the stimulus is readily apparent on multiple channels, including several where spontaneous neural activity was observed. (b) The same recording example as in (a), following high-pass filtering to isolate spiking activity.

We Figure 6a shows a set of traces for a single trial across the array on one of the recording sessions, before high-pass filtering is performed; Figure 6b shows the same data afiter high-pass filtering before spike detection. The raw, pre-cleaned voltage traces for this same trial is shown in Figure 1b, demonstrating that the vast majority of the artifact initially present has been removed. These high-pass filtered traces were then thresholded to extract spikes and then manually sorted into diferent units.

To evaluate the artifact removal process, we use prior knowledge about the multielectorde array channels. The most superficial (low numbered) and deepest (high numbered) channels were located above cortex and in white matter respectively, which we inferred from the signals observed on these channels, the density of neural signals on intermediate channels presumed to lie within cortex, and the known thickness of premotor cortex relative to the depth span of the probe. During non-stimulated trials, no detectable spiking waveforms were observed on these superficial and deep channels, therefore the presence of threshold crossings in the cleaned stimulation voltage traces on these channels would be surprising, and suggest that the cleaning procedure had failed to fully remove the stimulation artifact. However, as expected, the presence of evoked spiking activity during the stimulation window is limited to the intermediate channels, where spontaneous spiking activity was also observed (Figure 6). For closer inspection, we superimposed the raw voltage traces with cleaned, artifact-removed, high-pass filtered voltage traces for individual trials. Figure 7a-b shows such a comparison for the most superficial channel, where neither spontaneous nor evoked spiking activity was detectable. Figure 7c-d shows the same comparison for a channel with spontaneous spiking as well as robust evoked spiking activity.

**Figure 7.**
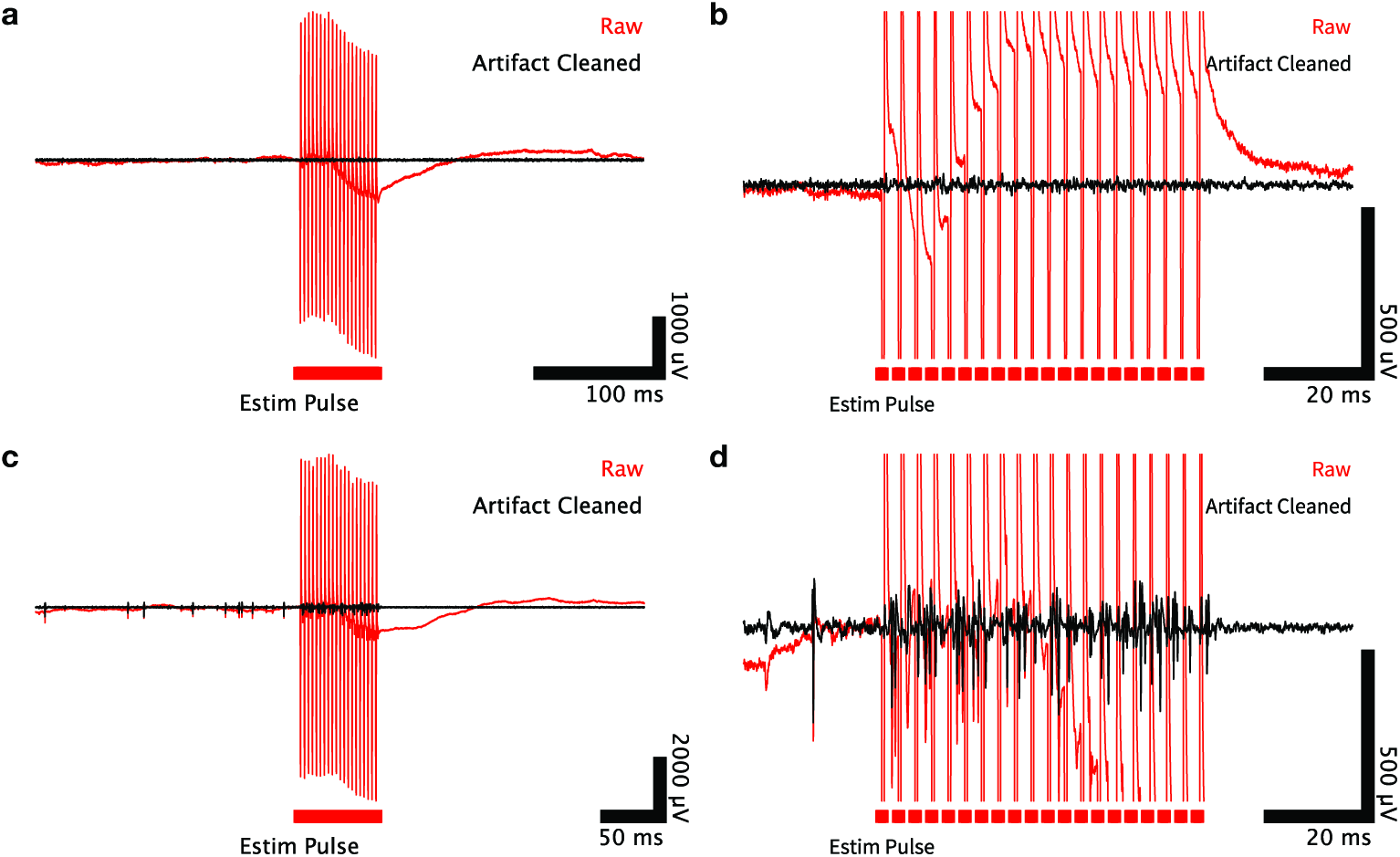
Comparison of raw and artifact cleaned signals. (a) Raw (red) and cleaned (black) voltage traces for a single trial on the most superficially located recording channel where no spontaneous activity was recorded, and no spiking activity is detected during stimulation. (b) Zoomed view of (a). (c) Same as (a) but for a channel located within cortex where spontaneous activity was present. (d) Zoomed view of (c) demonstrating evoked spikes visible during the stimulation period.

Secondly, it is essential to to ensure that threshold crossings detected during the stimulation period are likely due to real neural spiking activity, as opposed to residual transients due to stimulation artifact that persist through the cleaning procedure. We reasoned that threshold crossings created by residual artifact on a specific channel would not be expected to resemble the spontaneous spiking waveforms collected during non-stimulated trials on a per-trial basis. For each session, we generated a visual comparison of the spiking waveforms detected on non-stimulated trials with those detected within the stimulation window (putatively “evoked” spikes). These evoked spiking waveforms detected during the stimulation period were highly similar to the spiking waveforms detected during non-stimulation trials. Figure 8 demonstrates this similarity for a representative pair of experimental sessions. We note that these waveforms need not be identical, as evoked spiking activity from other neurons located further from the electrode could superimpose to corrupt the waveforms from nearby neurons, especially when this faraway spiking activity is highly synchronized by stimulation.

**Figure 8.**
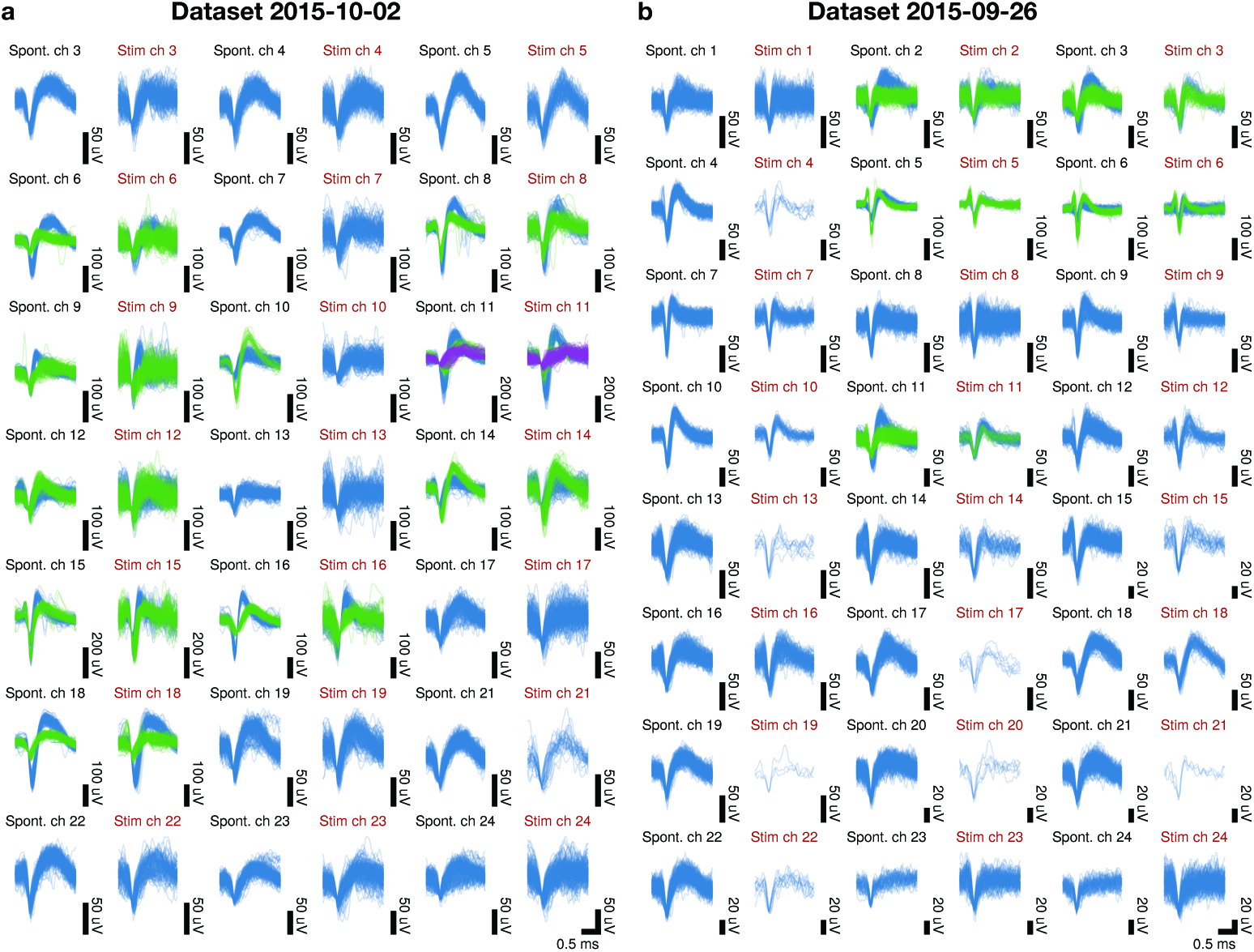
Recovered spiking waveforms detected during the stimulation period closely resemble spontaneously occurring waveforms recorded on the same channel. (a-b) Superimposed waveforms from each channel with spiking activity of two representative recording sessions. Each pair of columns compares spontaneously occurring (spont.) vs. peri-stimulus period (stim.) spiking waveforms. Waveform colors denote hand-sorted unit identity.

Lastly, we can quantify the amount of residual artifact directly by again utilizing channels that displayed no spontaneous spiking activity. Figure 9a compares the RMS voltage of these individual channels sampled from the post-cleaning stimulation window against a time window taken before stimulation. This metric captures the amount of baseline noise (largely thermal noise) on each channel, thereby providing an expectation for the variance of a given channel if the entirety of the corrupting artifact were successfully removed. For low amplitudes, the RMS voltage is ofiten slightly lower than the pre-stimualtion RMS, which is possible due to the greedy nature of the cleaning procedure. For larger amplitudes, the stimulation RMS remains acceptably close to pre-stimulation RMS. Figure 9b summarizes the similarity of spiking waveforms on each channel observed both spontaneously and during the stimulation window.

**Figure 9.**
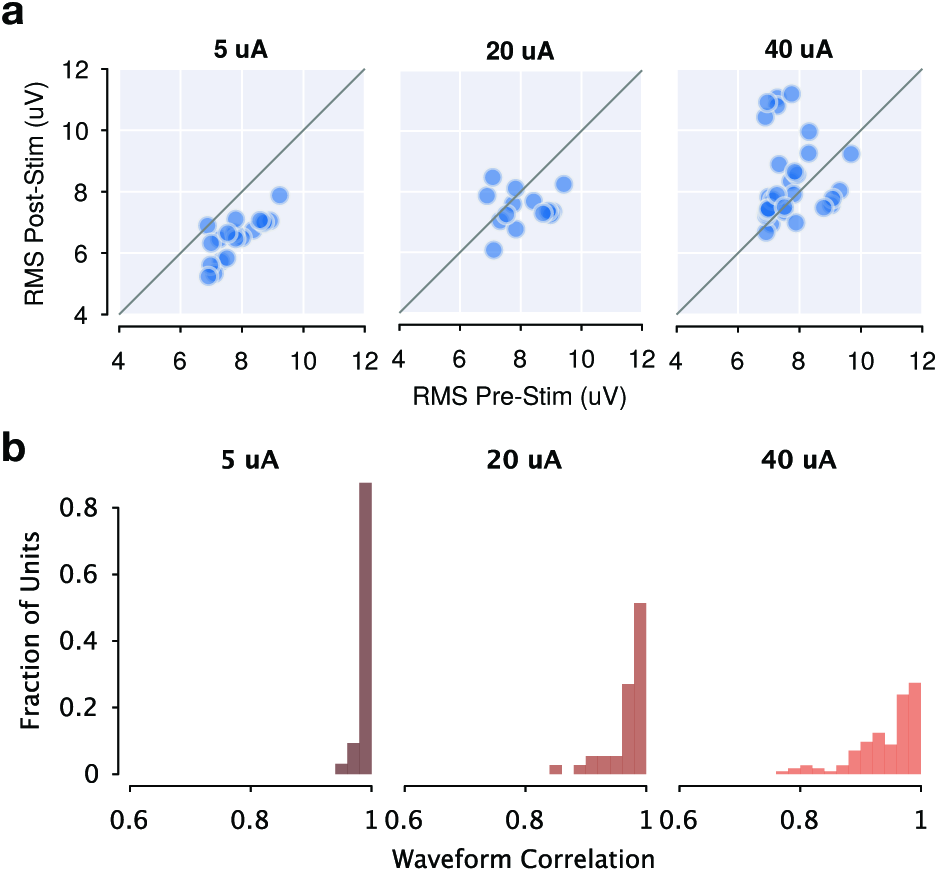
Summary of artifact removal quality across recording sessions. (a) RMS voltage on electrode channels which displayed no spontaneous spiking activity, across all recording sessions. Post-cleaning peri-stimulation RMS voltage is similar to spontaneous pre-stim RMS on each channel. (b) Summary of correlation coefficients between spontaneous spiking waveforms and recovered waveforms during stimulation, across all recovered units in all recording sessions.

## 4 Discussion

We discuss the main features of our method for estimating and removing electrical stimulation artifacts in comparison with existing approaches, as well as limitations of our approach and possible improvements. We also highlight a set of interesting neuroscientific arenas where the ability to observe electrically perturbed neural activity might be particularly illuminating, underscoring the utility of artifact removal methods.

### 4.1 Comparison to other methods

ERAASR exploits similarites in the electrical artifact observed across multiple channels, pulses within a stimulus train, and trials with repeated delivery of the same stimulus. Of these the shared structure of simultaneously recorded artifact across multiple channels was most useful. Most previously described attempts exploit only the common structure across trials, typically using an averaging or moving average approach (Hashimoto et al., 2002; Montgomery et al., 2005; Klink et al., 2017) to estimate artifacts on an individual channel. However, these approaches are necessarily quite sensitive to variability in artifact shape or amplitude across trials. While this variability may be small in relative terms, the much larger amplitude of artifacts relative to neural spiking waveforms can create meaningfully large errors in the residual signal, requiring that several milliseconds of signal in the vicinity of the stimulation pulse be discarded. Another common approach is to exploit differences in the shape of stimulation artifacts relative to spiking waveforms, by using curve-fitting algorithms to capture artifacts in the voltage signal but exclude spiking activity (Wagenaar and Potter, 2002; Erez et al., 2010). Similarly, these fits reliably capture artifact shape at a fixed delay from the stimulation pulse, but struggle to capture earlier portions of the transient (Erez et al., 2010), requiring peri-stimulus signal to be discarded. Motivated by the popularity of high-frequency pulse trains in ICMS experiemnts, our method addresses problem of observing neural activity through the entire stimulation pulse without discarding signal.

Our algorithm combines a series of simple, intuitive, linear operations along the channels, pulses, and trials axes of the data tensor. The core operation is principal components regression, in which PCA is used to identify a template describing common structure across a given axis of the tensor, followed by a regression step in which the artifact on a given channel (or pulse, or trial) is reconstructed by excluding that channel (pulse, trial) and its neighbors. This approach mitigates the risk of removing spiking signals during the cleaning process, exploiting the local nature of spiking responses across channels and the inherent biological variability in spiking responses on each pulse and each trial. Furthermore, each of these stages is intuitive and the results easily understood. A small set of key parameters, including the number of principal components and the set of neighbors excluded from the reconstruction at each stage, can be adjusted to trade-off between more aggressive cleaning and more veridical preservation of spiking waveform shape.

Our method operationally separates the process of artifact estimation from spike detection and extraction, which is performed as usual using typical high-pass filtering and thresholding on all voltage data afiter the cleaning process is complete. In contrast, a promising alternative method for multi-channel artifact removal is presented by Mena and colleagues (2016) for multielectrode array recordings of the retina. This approach employs a statistical model describing both the artifact and the spike generation process, jointly estimating artifact and spikes present in the voltage signals. This method exploits common structure of the artifact across a local group of channels around the stimulation source, but also requires that the electrical image of each neuron (the shape of spiking waveforms over several nearby electrodes) be known as an input to the cleaning algorithm. While this approach is highly effective and appropriate in the context of retinal recordings, where individual neurons are recorded on many densely spaced electrodes, the electrical images of neurons in primate cortex using typical multielectrode arrays are ofiten limited to one or two electrodes.

### 4.2 Limitations

First, our approach relies critically on the assumption that stimulation artifact linearly superimposes with neural spiking signals. Therefore, its utility is limited to stimulation configurations and amplitudes which occupy the linear, non-saturated regime of the amplifier. Practically, this sets a minimium distance between the electrode array and the stimulation source for a given stimulation amplitude. This distance could be designed such that the return path of the electrical current steers the electric field so as to minimize the recorded artifact amplitude (Rattay and Resatz, 2004). This problem can also be effectively managed in hardware, employing special circuitry to estimate and partially cancel artifacts online (Wichmann and Devergnas, 2011; Brown et al., 2008; Müller et al., 2012). A hybrid approach employing multi-channel artifact removal circuitry to prevent saturation using a predictable transformation of the recorded signal, supplemented with *post hoc* artifact cleaning procedure like the one described here could be effective across a much larger range of stimulation amplitudes.

Second, our algorithm sequentially removes shared structure across channels, pulses, and trials. We accomplish this by reshaping the voltage data tensor into a matrix (or set of matrices) so that familiar methods like PCA can be employed. This problem of finding shared structure naturally lends itself to tensor decomposition (Kolda and Bader, 2009), which could identify shared structure jointly over each of these axes and potentially improve the artifact estimation. Tensors naturally arise in neuroscientific data collection contexts, and decomposition methods are becoming increasingly useful for identifying population structure (Seely et al., 2016; Elsayed and Cunningham, 2017; Williams et al., 2017).

Lastly, our algorithm operates by greedily removing any shared structure as artifact, which can inadvertently remove or distort spiking waveform signals as well. In our dataset, we did not observe this while removing shared structure across channels, as we explicitly excluded adjacent channels from the reconstruction process. However, removing shared structure across pulses and trials assumes that spiking responses evoked by a stimulation pulse are variable in time. Moreover, with increasing stimulation amplitude, the temporal precision of evoked spikes may increase (Ranck, 1975; Butovas and Schwarz, 2003), which would lead to waveform distorion due to overjealous reconstruction and removal of spiking signals as artifact. We experimented with using matched-filters for robust spike detection while relaxing the artifact estimation technique (data not shown), though we did not explore whether this would be applicable at higher currents and more synchronous evoked spiking. In these circumstances, we expect that a joint estimation of artifact and spiking as proposed by Mena et al. (2016) could be adapted to more effectively recover highly regular evoked spiking by exploiting other differences in the structure of spikes from artifact.

### 4.3 Motivation for direct observation of electrically perturbed neural activity

ICMS is a well-established and popular technique for perturbing neural activity and probing the causal contributions of a certain brain region to specific cognitive functions (Cohen and Newsome, 2004; Histed et al., 2013; Clark et al., 2011; Tehovnik, 1996). Here we argue that recent results suggest that the effect of electrical stimulation on neural activity may be significantly more complex than previously realized, which implies that direct observations of the net effect on neural activity in the perturbed region as well as upstream and downstream areas could reveal the precise mode by which microstimulation modulates behavior.

The initial conception of ICMS is that stimulation activates most neurons within a sphere surrounding the electrode tip (Stoney et al., 1968; Tehovnik, 1996; Tehovnik et al., 2006). This idea derives originally from the findings of Stoney et al. (1968), who cleverly side-stepped the issue of stimulation artifacts by using collisions of antidromic and orthodromic spikes in a dual-stimulation paradigm as an indirect measure of local neuronal activation. Using this technique, they estimated that 10μA and 100 μA ICMS currents would activate most pyramidal cells in a local ball extending 100 μm and 450 μm in radius, respectively. A wealth of additional research has also attempted to carefully characterize the sensitivity of various elements of the CNS to stimulation, as a function of current amplitude, pulse duration and shape, electrode configuration, etc. (e.g. Ranck, 1975; Asanuma et al., 1976; Tehovnik, 1996; Rattay, 1999; Tehovnik and Slocum, 2007; Marcus et al., 1979; Nowak and Bullier, 1996; Nowak and Bullier, 1998b; Nowak and Bullier, 1998a; Kimmel and Moore, 2007). The primary findings of this body of work are that stimulation primarily evokes spikes at axons, in particular the excitable axon initial segment, and that the likelihood of evoking a spike in a neuron depends on distance, pulse duration, and current according to a relatively simple power law. The spatial profile of ICMS has been bolstered by multiple behavioral studies as well (e.g. Murasugi et al., 1993; Tehovnik et al., 2004; Tehovnik et al., 2005; Bartlett et al., 2005; Tehovnik and Sommer, 1997). These studies derive an estimate of the effective activation volume and dependency on stimulation parameters by combining behavioral readouts with known features of cortical spatial organization. For example, Murasugi et al. (1993) found that low current ICMS delivered to monkey area MT could bias perceived dot motion direction, but that ICMS currents above 20μA less effectively biased perception. The authors conclude that below this threshold current, electrical activation is confined to a single “directional” column in MT, about 0.2mm in diameter (Albright et al., 1984).

However, recent evidence challenges this traditional view of localized, dense activation. Using two-photon calcium imaging, Histed et al. (2009) demonstrated that low-intensity ICMS activated a sparse collection of neurons distributed in a relatively large volume extending several hundred μm. In two experiments in cats, the authors also found neurons as far as 4mm away activated by 10μA stimulation, perhaps resulting from lateral axons extending multiple millimeters within layer 2/3 of visual cortex in cats but not rodents. The widely distributed collection of neurons is likely activated by antidromic stimulation of their axons. By advancing the electrode 30 μm, the authors found that an entirely non-overlapping sparse set of cells was activated, suggesting that ICMS at low currents (10-25 μA) activates axonal processes within ≈15 μm of the electrode tip. At higher currents, more cells within the distributed volume were activated, though all experiments were conducted with currents ≤30 μA.

Additionally, many mapping studies have focused on direct activation in response to a single isolated current pulse. Nonetheless, it has long been known that ICMS induces transsynaptic spiking in connected neurons (Stoney et al., 1968; Butovas and Schwarz, 2003). As larger currents activate more neurons, the effect of transsynaptic activation may be enhanced due to the summation of many from this larger directly activated population. Cortically evoked spiking may also depend on subthreshold activity in neurons, which is continuously modulated by ongoing task or stimulus-evoked responses (Kara et al., 2002). When multiple pulses are delivered, as is typical in experiments applying ICMS in behaving animals, short-term synaptic depression commonly seen in cortex would also modulate the effectiveness of transynaptically evoked spiking (Thomson and Deuchars, 1994; Stratford et al., 1996; Deisz and Prince, 1989; Varela et al., 1997). When longer pulse trains are used, particularly at high currents, electrical stimulation might readily engage a large collection of neural circuits while driving temporally complex neural responses (Strick, 2002; Graziano et al., 2002).

It is also well-established that larger stimulation currents can recruit inhibitory interneurons and result in inhibition of cortical responses lasting hundreds of milliseconds afiter stimulation (Berman et al., 1991; Watanabe et al., 1966; Li et al., 1960; Chung and Ferster, 1998; Creutzfeldt et al., 1966; Kara et al., 2002; Masse and Cook, 2010; Seidemann et al., 2002; Butovas et al., 2006; Butovas and Schwarz, 2003). Additionally, in an experiment combining cortical stimulation with fMRI and extracellular recording, Logothetis et al. (2010) demonstrated that high-frequency ICMS of the thalamic lateral geniculate nucleus (LGN) initially excites (but later suppresses) monosynaptically connected primary visual cortex (V1), but suppresses responses in downstream extrastriate cortices. This suppression was abolished by bicuculine infusion. The authors concluded that, high-frequency ICMS pulse trains disrupt normal information flow along cortico-cortical pathways, by recruiting strong inhibition that prevents the spread of activation beyond the regions whose direct aferents were stimulated.

Taken together, the net effects of ICMS on the firing patterns of cortical neuronal populations may be quite complicated. The directly activated population is likely sparse, widely distributed, and stochastic due to a strong dependence on axonal proximity to the stimulating electrode. Moreover, the transsynaptically activated population of cells undergoes a complex, time-varying pattern of modulation that reflects circuit micro-architecture, local circuit dynamics, short-term synaptic plasticity, recruitment of inhibition, and an interaction with ongoing task-related activity. With this evident complexity, we feel that arriving at an accurate interpretation of the behavioral consequences of electrical microstimulation first requires careful measurement of its effects on local neuronal population activity (Jazayeri and Afraz, 2017; Otchy et al., 2015). Furthermore, an accurate picture of electrically modulated neural activity could elucidate the circuit mechanisms underlying intriguing aspects of the interaction between microstimulation and behavior, such as how electrically evoked eye movements interact with visual guided eye movements (Sparks and Mays, 1983), how precise timing of electrical pulses modulates saccadic effects (Kimmel and Moore, 2007), how perceptural decisions are biased by stimulation of LIP (Hanks et al., 2006), how motor preparation recovers subsequent to electrical disruption (Churchland and Shenoy, 2007; Shenoy et al., 2011), and how motor cortical stimulation can evoke complex movements converging on a specific endpoint (Graziano et al., 2011) and effectively nullify the contribution of goal-directed behavior in the motor cortical output (Griffin et al., 2011). A detailed comparison with the effect of optogenetic stimulation could clarify the differences (Gerits and Vandufel, 2013; Diester et al., 2011; Ohayon et al., 2013) and similarities (Dai et al., 2013) observed between the two modalities’ effects on behavior. Lastly, a detailed model of microstimulation’s interaction with cortical dynamics could facilitate delivery of more reliable or realistic artificial sensory perception in visual, auditory, and motor prostheses (Otto et al., 2005; Tehovnik et al., 2009; O’Doherty et al., 2011; Bensmaia and Miller, 2014; Dadarlat et al., 2015).

## 5 Conclusion

We developed ERAASR, an algorithm for estimating and removing artifacts caused by electrical stimulation on multielectrode array experiment. ERAASR assumes that artifact is shared over many channels and that evoked transients are highly repeatable across pulses and trials, whereas spiking activity is highly local and temporally jittered. Shared structure in the voltage signals is identified and removed sequentially across channels, across pulses in a stimulus train, and across trials, using straightforward linear methods. We believe our technique will be useful to neuroscientists in drawing precise causal links between perturbations and the effect of stimulation on neural activity and behavior and aid in the design and implementation of enhanced neural prosthetic systems capable of restoring lost sensation and facilitating precise control.

## 6 Acknowledgments

We thank Tirin Moore, Rob Franklin, and Paul Venable for helpful discussions. This article was supported by the following grants: Christopher and Dana Reeve Paralysis Foundation, Burroughs Welcome Fund Career Awards in the Biomedical Sciences„ Defense Advanced Research Projects Agency Reorganization and Plasticity to Accelerate Injury Recovery N66001-10-C-2010, US National Institutes of Health Institute of Neurological Disorders and Stroke Transformative Research Award R01NS076460, US National Institutes of Health Director’s Pioneer Award 8DP1HD075623-04, US National Institutes of Health Director’s Transformative Research Award (TR01) from the NIMH 5R01MH09964703, Defense Advanced Research Projects Agency NeuroFAST award from BTO W911NF-14-2-0013, Howard Hughes Medical Institute, NSF GRFP, and NSF IGERT.

